# Spatio-temporal modeling of the effect of selected environmental and land-use drivers on acid grassland vegetation

**DOI:** 10.1101/2020.10.25.353839

**Authors:** Christian Damgaard

**Affiliations:** Department of Bioscience, Aarhus University, Vejlsøvej 25, 8600 Silkeborg, Denmark

**Keywords:** Joint distribution of plant abundance, spatial and temporal variation of plant cover, hierarchical Bayesian models, pin-point cover data, structural equation modelling, acid grassland vegetation

## Abstract

Spatial and temporal pin-point plant cover monitoring data are fitted in a structural equation model in order to understand and quantify the effect of selected environmental and land-use drivers on the observed variation and changes in the vegetation of acid grasslands. The important sources of measurement- and sampling uncertainties have been included using a hierarchical model structure. Furthermore, the measurement- and sampling uncertainties are separated from the process uncertainty, which is important when generating ecological predictions. Generally, increasing atmospheric nitrogen deposition leads to more grass-dominated acid grassland habitats at the expense of the cover of forbs. Sandy soils are relatively more acidic, and the effect of soil type on the vegetation includes both direct effects of soil type and indirect effects mediated by the effect of soil type on soil pH. Both soil type and soil pH affected the vegetation of acid grasslands. Even though only a small part of the temporal variation in cover was explained by the model, it will still be useful to quantify the uncertainties when using the model for generating local ecological predictions and adaptive management plans.

## Introduction

Acid grasslands in Atlantic lowland areas are semi-natural habitats often found on oligotrophic more or less acidic sandy soils (Galvánek and Janák 2008). Acid grasslands are low productivity grasslands, which persist due to extensive grazing and occasional mowing, and they are threatened both by agricultural intensification with nutrient addition and increased livestock densities as well as by land abandonment, where grasslands are overgrown by shrubs and trees (Galvánek and Janák 2008; Timmermann et al. 2015). Furthermore, atmospheric nitrogen deposition has been shown to be associated with changes in species composition and a reduction in plant species richness of especially forbs (Damgaard et al. 2011; Helsen et al. 2014; Stevens et al. 2011; Stevens et al. 2006; Stevens et al. 2010). Additionally, changes in precipitation patterns due to climate changes may cause modifications in interspecific plant competitive interactions and alterations in plant community compositions (Galvánek and Janák 2008; Herben et al. 2003; Hopkins and Del Prado 2007).

Most large-scale studies of the effect of anthropogenic environmental and land-use changes on acid grassland vegetation have been conducted using spatial regression methods on static data. However, in order to get closer to the underlying causes of the observed vegetation changes, it is preferable to study both spatial and temporal variation (Damgaard 2019a). Here, spatial and temporal pin-point plant cover data, which are collected as part of a Danish ecological monitoring program (Nielsen et al. 2012), are fitted to a spatio-temporal structural equation model (SEM) in order to understand and quantify the effect of environmental and land-use drivers on the observed variation and changes in the vegetation of acid grasslands. Furthermore, latent geographic factors of the large-scale spatial variation will be estimated (Ovaskainen et al. 2016).

Pin-point cover data are readily aggregated into species classes, and in order to simplify the model and generalize the results, all the observed higher plant species are divided into four species classes: grass, other graminoids, dwarf shrubs and other forbs. This species classification scheme is motivated by important differences in plant functional types as well as conceivable nature conservation issues of acid grasslands. The joint distribution of the multivariate cover data of the four species classes are analyzed using a reparametrized Dirichlet – multinomial distribution, where the typically observed spatial aggregation of plant species is taken into account (Damgaard 2015; 2018).

When fitting ecological and environmental models to data, there are often important sources of measurement- and sampling uncertainties, and it may be important to model this uncertainty to avoid model- and prediction bias (Damgaard 2020b). Consequently, the applied SEM has been fitted within a Bayesian hierarchical model structure using latent variables, which have the added advantage that the modelled measurement- and sampling uncertainties are separated from the process uncertainty. The hierarchical SEM approach and the motivation for using it are explained in more detail in Damgaard (2019b), where a similar model was fitted to ecological monitoring data from wet heathlands.

The aim of the present study is to model the effect of selected environmental drivers; soil type, pH, nitrogen deposition, precipitation and grazing on the large-scale spatial distribution and temporal changes of the acid grassland vegetation. The obtained model results will be discussed in the light of the abovementioned threats to acid grassland habitats.

## Materials and Methods

### Sampling design

Hierarchical time series data from 105 acid grassland sites (Fig. 1) that had been monitored at least three times in the period from 2007 to 2014 were used in the analysis. All sites included several plots classified as acid grassland (EU habitat type: 6230) according to the habitat classification system used for the European Habitat Directive (EU 2003). The area of the sites ranged from 0.77 ha to 382 ha, with a median area of 12.6 ha. A total of 1135 unique plots were used in the analysis. The sampling intensity was irregular among sites and years, but typically between ten and forty plots were sampled from each site at each time point. Including resampling over the years, a total of 3444 plot data were used in the analyses. The plots were resampled with GPS-certainty (< 10 meters).

**Fig. 1.**
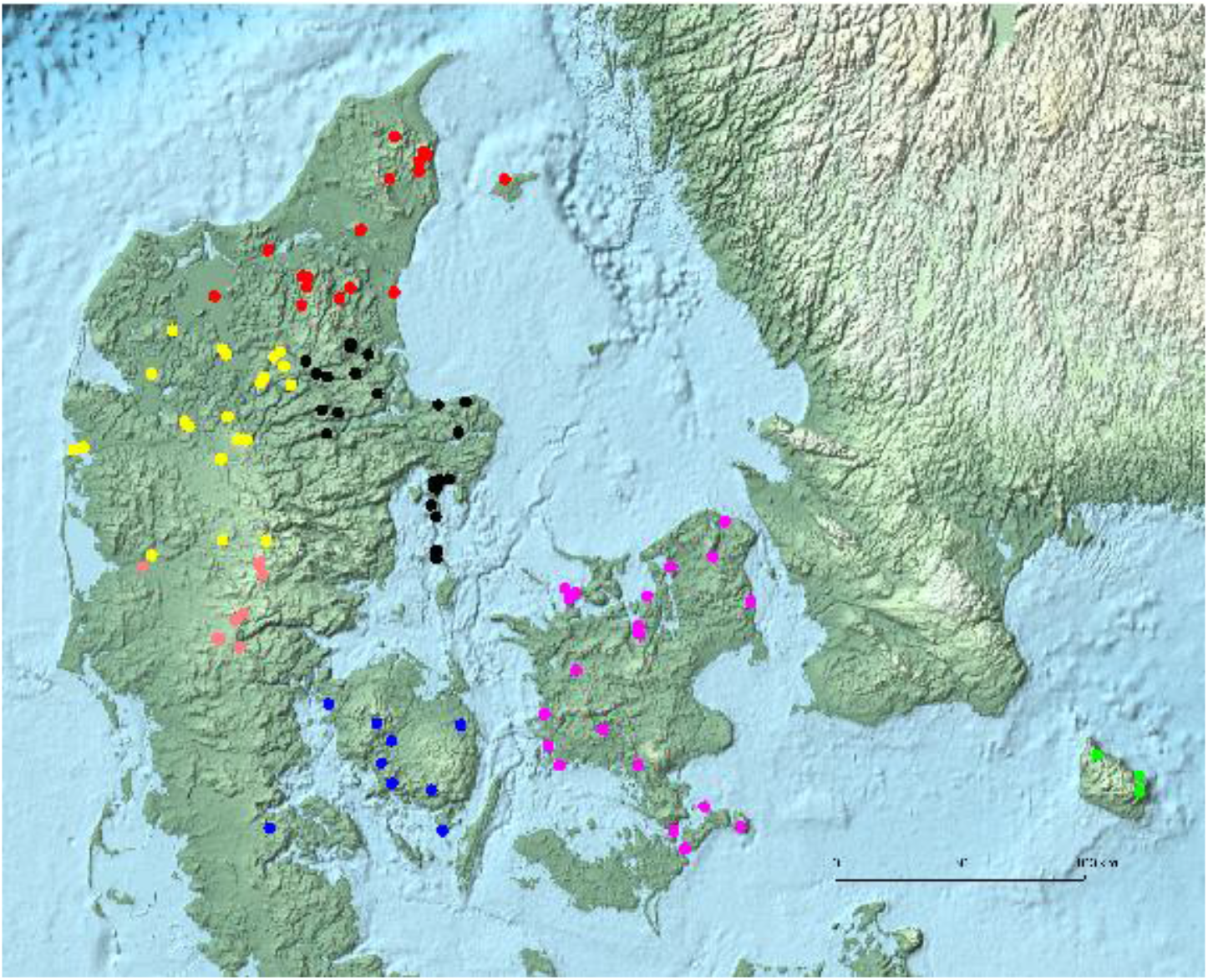
Map of the 105 Danish acid grassland sites. The different colors represent a classification of the different sites into seven geographical regions.

The data are a subset of the data collected in the Danish habitat surveillance program NOVANA (Nielsen et al. 2012; Nygaard et al. 2016).

### Variables and measurements

#### Plant cover data

The plant cover, which is the relative projected area covered by a species, was measured for all higher plants by the pin-point method (Damgaard and Irvine 2019; Levy and Madden 1933; Lindquist 1931) using a square frame (50 cm X 50 cm) of 16 grid points that were equally spaced by 10 cm (Nielsen et al. 2012). Cover data from the 25 most common species in Danish acid grassland are summarized in Table S1.

Since the pin-point cover data were recorded for each pin separately, the species cover data are readily aggregated into cover data for classes of species at a higher taxonomic or functional level. If a pin has hit two different plant species from the same species class, then the two hits are counted as a single hit of the species class. In this study, the observed higher plant species were divided into four species classes: grass species *(Poaceae),* other graminoids *(Cyperaceae* and *Juncaceae),* dwarf shrubs *(Ericaceae)* and other forbs (all higher plant species, except graminoids, *Ericaceae,* trees and bushes). The mean site covers of the four species classes are shown in fig. 2.

**Fig. 2.**
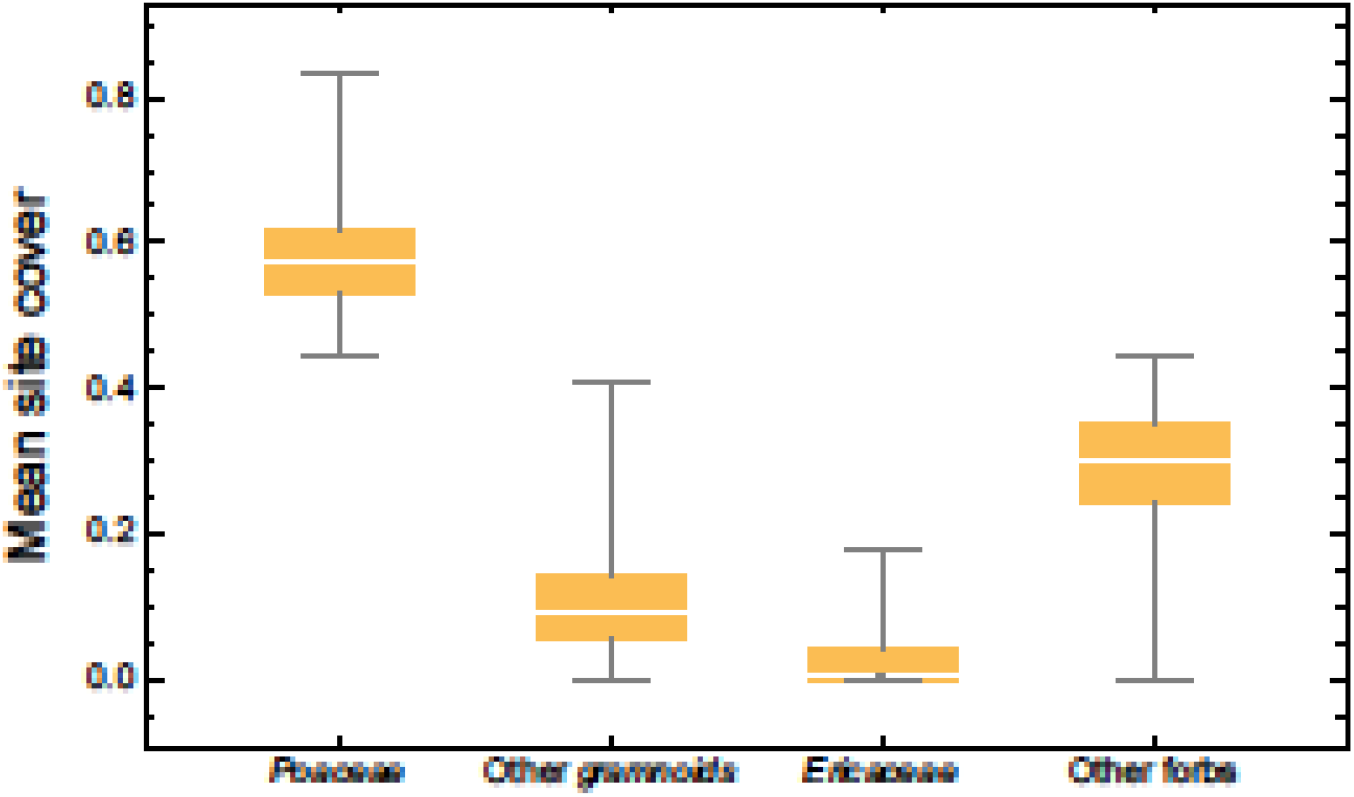
Box plots of the mean site cover of the four species classses.

#### Nitrogen deposition

Nitrogen deposition at each plot was calculated for each year using a spatial atmospheric deposition model in the period from 2005 to 2014 (Ellermann et al. 2012). The mean site nitrogen deposition ranged from 5.66 kg N ha^-1^ year to 21.92 kg N ha^-1^ year^-1^, with a mean deposition of 12.87 kg N ha^-1^ year^-1^.

#### PH

Soil pH was measured in randomly selected plots from the uppermost 5 cm of the soil (four samples were amassed into a single sample). The soil sampling intensity was irregular among sites and years, but typically between one and four plots were sampled from each site at each time point. When a plot was resampled, the pH at the plot was calculated as the mean of the samples. In total, 632 independent soil pH values were used in the analysis. The soils were passed through a 2mm sieve to remove gravel and coarse plant material, and PH_KCI_ was measured on a 1 M KCl-soil paste (1:1). The measured soil pH ranged from 2.7 to 7.5, with a mean soil pH of 4.36.

#### Soil type

The texture of the top soil for each site was obtained from a raster based map of Danish top soils (Greve et al. 2007). The classification of the soil (JB-nr.) is on an ordinal scale with 1: coarse sandy soil, 2: fine sand soil, 3: coarse loamy soil, 4: fine loamy soil, 5: clay soil. There were some records with other soil types, but because of possible classification errors they were treated as missing values. The mean soil type was 2.56.

#### Precipitation

Site-specific precipitation was measured by the average annual precipitation in the period 2001 to 2010, with a spatial resolution of 10 km (DMI 2014). The annual precipitation ranged from 587 mm to 932 mm, with a mean precipitation of 736 mm.

#### Grazing

Signs of grazing, e.g. the presence of livestock or short vegetation within fences, was recorded by the observer at each plot for each year since 2007 as a binary variable (sign of grazing = 1, no sign of grazing = 0). The mean grazing at the site level was 0.72.

#### Geographic regions

The 105 acid grassland sites were initially grouped within geographic squares with an edge of 50 km. This initial grouping was modified by aggregating neighboring groups, so that each geographic group consisted of at least two sites. This resulted in seven geographic regions (Fig. 1) that were used to investigate possible latent geographic factors (Ovaskainen et al. 2016).

### Structural equation model

The variables were modelled in a SEM based on current knowledge on the causal effect relationships among the studied abiotic variables and grazing and their effect on the vegetation in acid grasslands (Ejrnæs et al. 2009; Galvánek and Janák 2008; Stevens et al. 2006; Williams and Anderson 1999). The SEM was fitted within a Bayesian hierarchical framework with structural equations and measurement equations in an acyclic directed graph (Fig. 3), where each arrow is modelled by a conditional likelihood function (Damgaard 2019b).

**Fig. 3.**
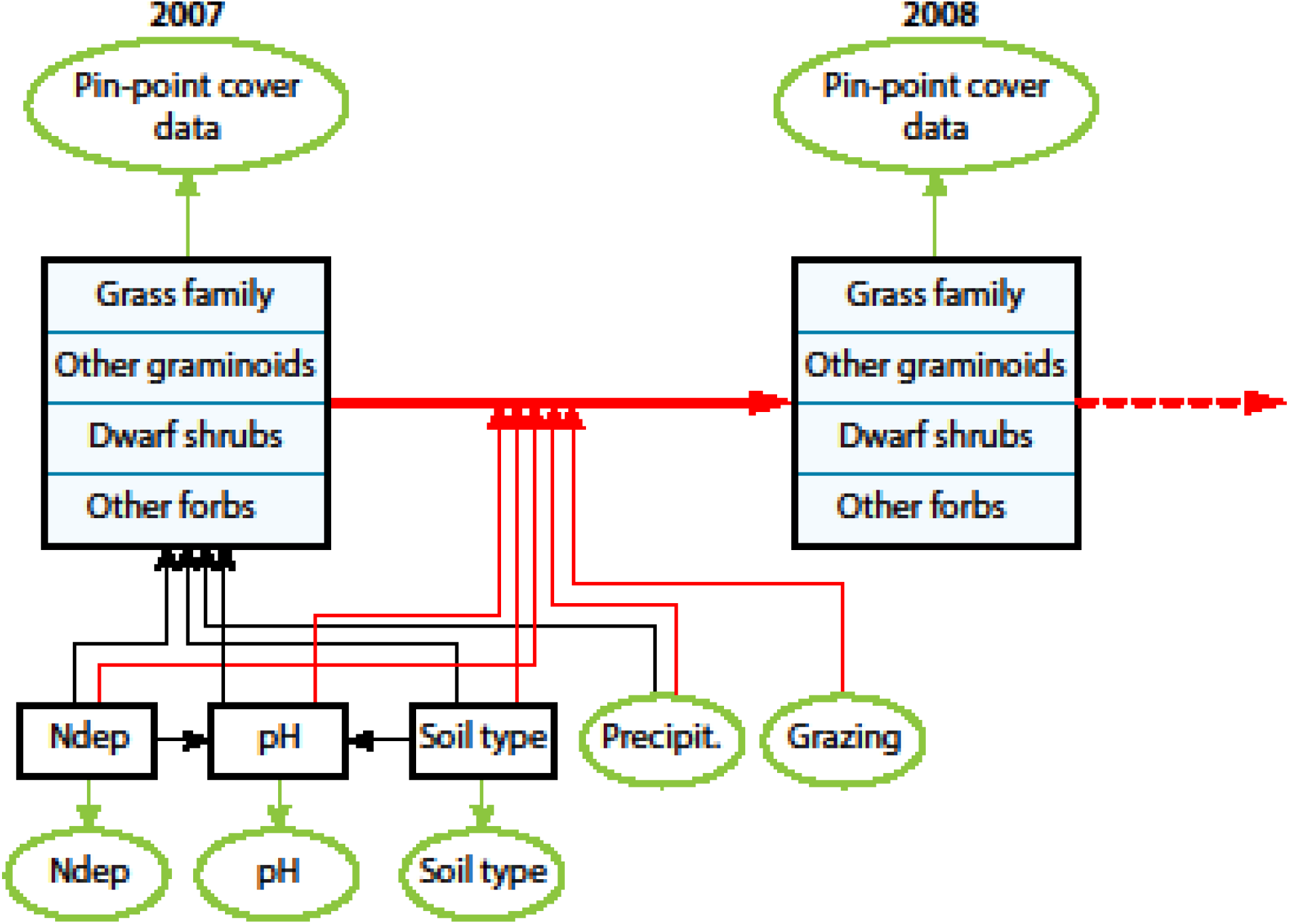
Outline of SEM. The spatial variation in vegetation cover in 2007 is modelled by nitrogen deposition (Ndep), soil pH (pH), soil type and precipitation (Precipit.). The yearly change in vegetation cover from 2007 to 2014 (only a single yearly change is shown in the figure) is modelled by all the former variables as well as grazing. The black boxes are latent variables and the green ovals are data. The black arrows denote spatial processes, and the red arrows denote temporal processes.

#### Structural equations

The structural equations consist of a mixture of measured data and latent variables, which are the true, but unknown, mean of the underlying data (Fig. 3). The latent variables are denoted by upper case Latin letters, data are denoted by lower case Latin letters, and model parameters are denoted by Greek letters.

The relative cover of the three species, *s*: grass, other graminoids, and dwarf shrubs at each site, *i,* for the first year (1 = 2007) and for all subsequent years, *y,* where cover was measured, are modeled as latent variables, *Q_s,i,y_*. Furthermore, the latent geographic factor *(Geo_i_*) and the true, but unknown, mean of nitrogen deposition *(N_i_*), soil pH (*R_i_*), and soil type (*S_i_*) were modeled by site-specific latent variables. Site-specific average precipitation is denoted by *p_i_*.

The large-scale spatial variation in the relative cover the first year is modelled as,

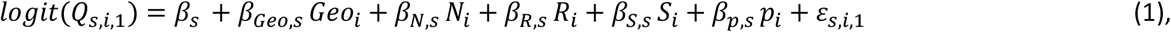

where the structural uncertainty in the spatial variation of the logit-relative covers transformed 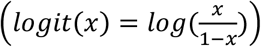 is assumed to be normally distributed, 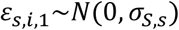.

The yearly change in the logit-transformed relative cover in subsequent years is modelled as,

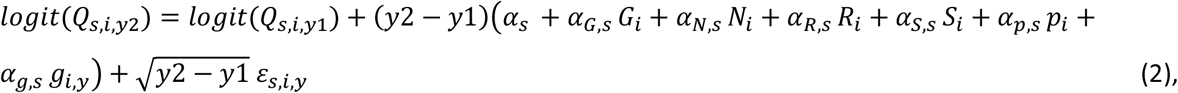

where (*y*2 — *y*1) is the years between successive sampling at site *i, G_i_* is the latent variable that measures grazing, and the structural uncertainty in the temporal variation of the logit-transformed relative covers are assumed to be normally distributed, *ε_s,i,y_*~*N*(0, *σ_T,s_*).

The site-specific soil pH in the top soil was modelled as,

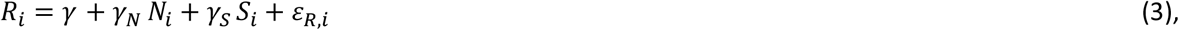

where the structural uncertainty in soil pH is assumed to be normally distributed, *ε_R,i_*~*N*(0,*σ_SR_*).

The latent geographic factors were assumed to come from a Gaussian process, *G~N*(0,1) (Haran 2011).

#### Measurement equations

The measurement equations link the latent variables to the measured or model-calculated data with conditional likelihood functions (Fig. 3):

The observed pin-point cover data of grass, other graminoids, dwarf shrubs and other forb at site *i*, plot *j* and year y, *q_s,i,j,y_* were assumed to be jointly distributed according to a Dirichlet – multinomial mixture distribution, *q_s,i,j,y_*~*DM*(*Q_s,i,y_, δ*), with *Q_s,i,y_* as the relative mean cover of grass, other graminoids and dwarf shrubs at site *i,* and small-scale spatial aggregation parameter *δ* (Damgaard 2015; 2018). If the plot had been sampled before then within-plot auto-correlation was taken into account by, *q_s,i,j,y_*~*DM*(*Q_s,i,y_* + *ρ_s_*(*q_s,i,j,yp_* – *Q_s,i,yp_*), *δ*), where *ρ_s_* are correlation coefficients and *yp* denote the year of the pervious sample at the plot (Damgaard 2012).

The average model calculated nitrogen deposition at site *i* and year *y,n_i,y_*, was assumed to be normally distributed, *n_i,y_*~*N*(*N_i_, σ_N_*), where *σ_N_* is the yearly variation in model calculated nitrogen deposition.

The measured soil pH at site *i* and plot *j, r_i,j_*, was assumed to be normally distributed, *r_i,j_*~*N*(*R_i_, σ_R_*), where *σ_R_* is the within site variation in soil pH. At three sites there were no measurements of soil pH, and at these sites *R_i_* was not conditioned by measured data, but only by the structure of the fitted SEM (Fig. 3).

The predicted texture of the top soil at site *i* and plot *j, S_i,j_* was assumed to be normally distributed, *s_i,j_*~*N*(*S_i_, σ_s_*), where *σ_s_* is the within site variation in soil type.

### Estimation and statistical inference

The SEM was parameterized using numerical Bayesian methods. The integrated likelihood function of the SEM model was found by multiplying the conditional likelihood functions specified above and outlined in figure 3, using a first order Markov assumption (Clark 2007). The joint posterior distribution of the parameters and the latent variables were calculated using Markov Chain Monte Carlo (MCMC), Metropolis-Hastings, simulations with a multivariate normal candidate distribution. The MCMC had 120,000 iterations, with a burn-in period of 70,000 iterations.

The prior distributions of all parameters were assumed to be uniformly distributed either as improper priors or in their specified domain, except standard deviation parameters of the normal distribution that were assumed to be inverse gamma distributed, (*σ_x_*~*IG*(0.1,0.1). The prior distributions of the cover latent variables were assumed to be uniformly distributed within 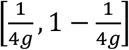, where *g* is the number of pins in a pin-point frame (Damgaard 2012). The other latent variables were assumed to be improper uniformly distributed.

Plots of the sampling chains of all parameters and latent variables were inspected to check the mixing properties of the used sampling procedure. Additionally, the overall fitting properties of the model were checked by inspecting the regularity and shape of the marginal distribution of parameters as well as the distribution of the deviance (= —2 *log L*(*Y|Θ*)). The efficiency of the MCMC procedure was assessed by inspecting the evolution in the deviance.

In order to check the fit of the model, the mean latent logit-transformed cover variables were plotted against the expected logit-transformed cover for both the spatial and temporal structural processes. Furthermore, the Dunn-Smyth residuals (Dunn and Smyth 1996) of the marginal observed cover data of the four species classes were calculated with the mean latent cover variables as the expected covers *(q_i_*) and the mean small-scale spatial aggregation parameter *(δ)* (Damgaard 2018).

Statistical inferences on the parameters were based on the estimated marginal posterior distribution of the parameters. Standardized regression coefficients (path coefficients) were calculated by multiplying the estimated partial regression coefficients with the standard deviation of the independent latent variable and dividing with the standard deviation of the dependent latent variable.

All calculations were done using *Mathematica* version 11.0.1 (Wolfram 2016), and the *Mathematica* notebook with additional diagnostic plots are provided as an electronic supplement.

## Results

There was considerable covariation among the studied abiotic variables at the 105 acid grassland sites (Fig. 4), which is expected to lead to covariance among parameter estimates (Table S2) and therefore affect the fitting properties of the model negatively. Plots of the mean latent vs. expected logit-transformed cover variables demonstrated an acceptable fit of the spatial variation in cover (Fig. 5A; 55% of the variation is explained), whereas the model fitted the temporal process of the change of cover poorly (Fig. 5B; only 4% of the variation is explained). Generally, the distribution of the Dunn–Smyth residuals of the marginal observed cover data of the four species classes were approximately normally distributed (Fig. S1).

**Fig. 4.**
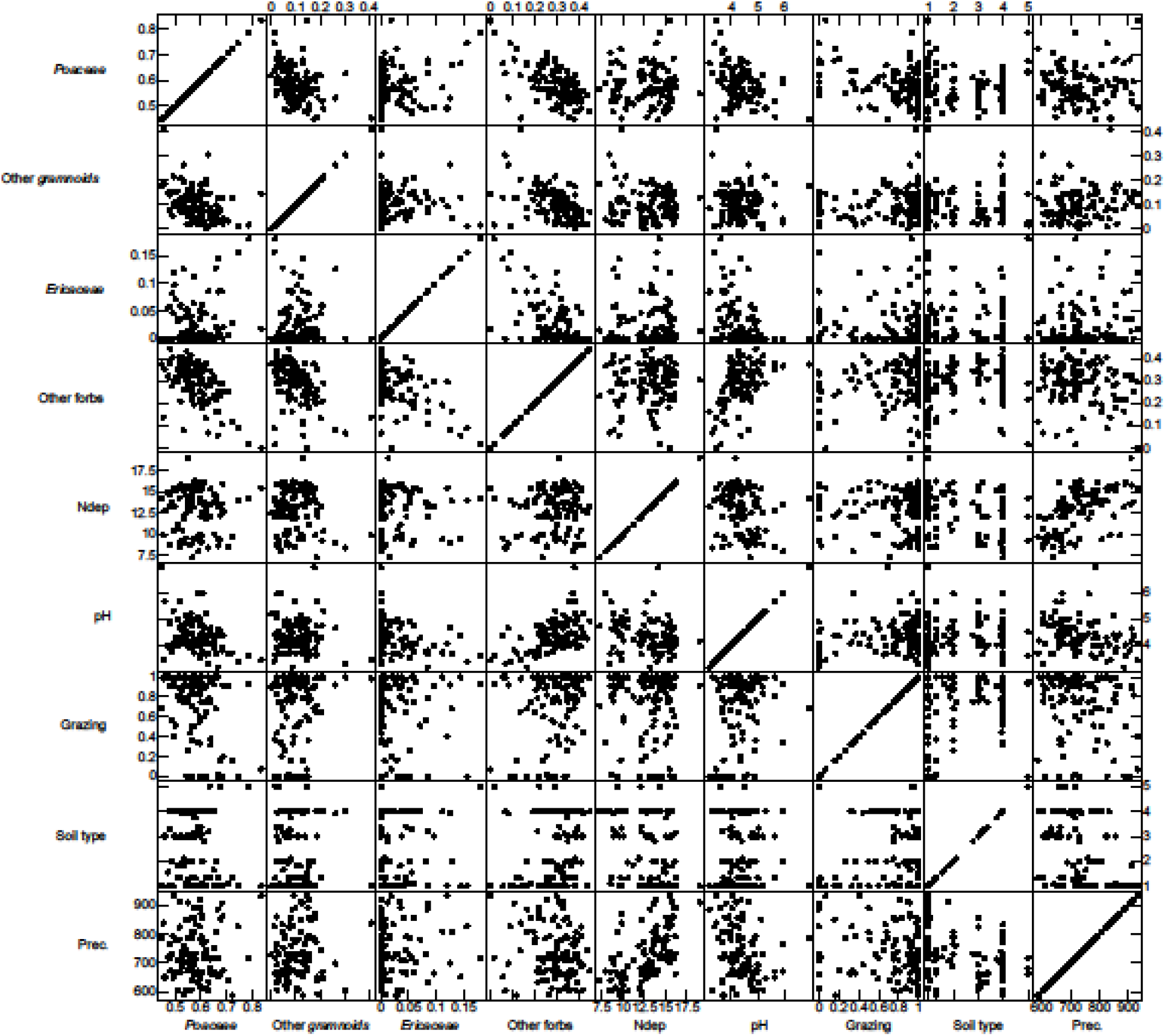
Pairwise scatter plot of the mean at site level for all variables.

**Fig. 5.**
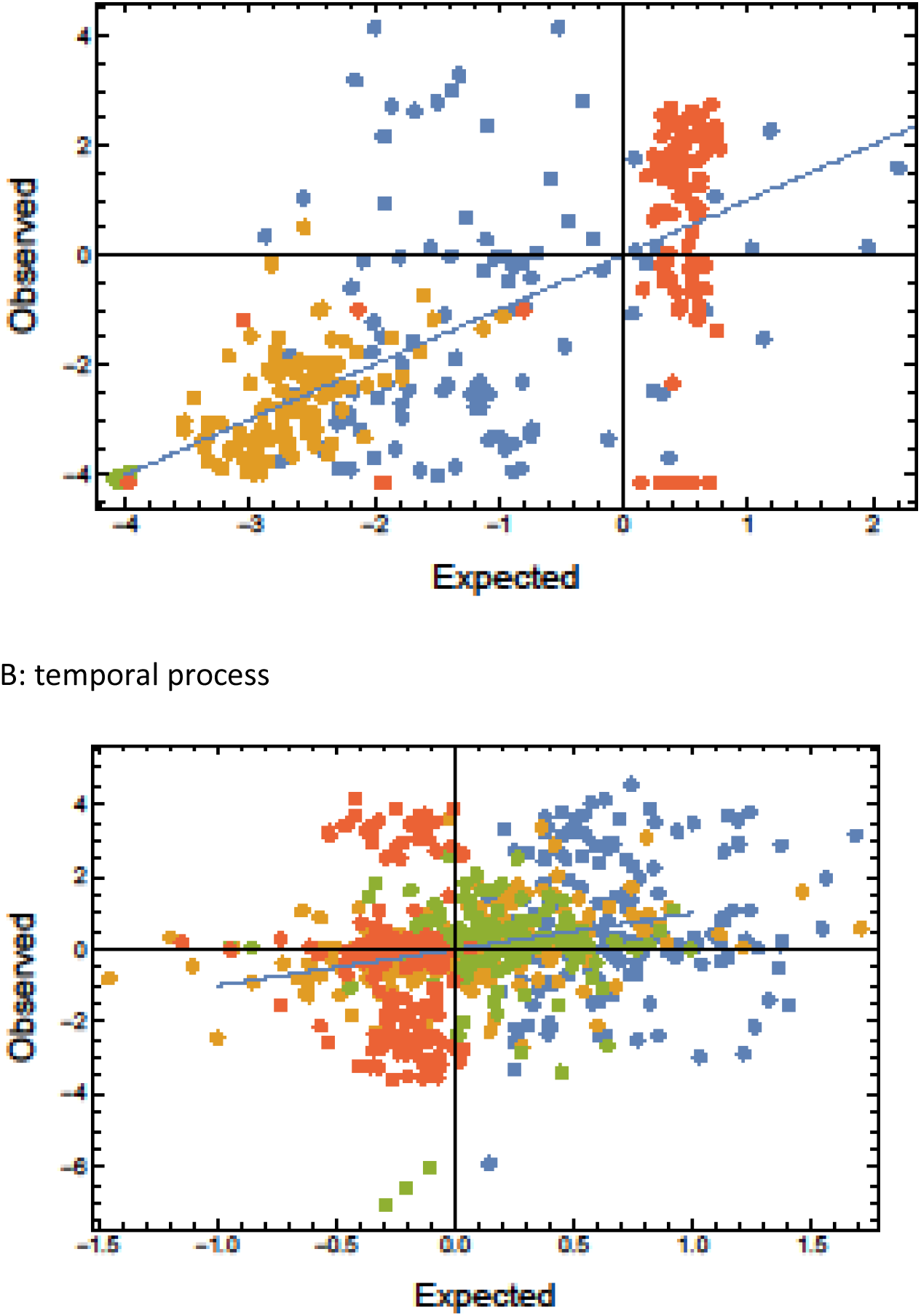
Plots of observed vs. expected of initial logit-transformed cover (A) and subsequent change of logit-transformed cover (B). Blue: grass, yellow: other graminoids, green: dwarf shrubs, and red: other forbs.

The statistical modelling uncertainty is partitioned into measurement uncertainty of plant cover, nitrogen deposition, soil pH and soil type and structural uncertainties due to the modelled large-scale spatial variation and temporal processes. The most important source of measurement uncertainty was the plant cover measurement due to the significant small-scale spatial aggregation of plant species, which were modelled by the Dirichlet – multinomial mixture distribution and, more specifically, by the parameter *δ.* The median estimated value of *δ* was 0.18 with a relatively small credible interval (Table 1). This estimated amount of spatial aggregation significantly increased the measurement uncertainty of the expected cover data compared to the case of randomly distributed plant species. If this over-dispersion of the pin-point cover data relative to the random expectation is not taken into account in the statistical model, then the signal to noise ratio will be severely upward biased and may lead to biased model results. Generally, the structural uncertainties of the model are rather high, except for the large-scale spatial variation of dwarf shrubs, which is one-order of magnitude lower than the other estimated structural uncertainties (Table 1, Fig. 5).

**Table 1:**
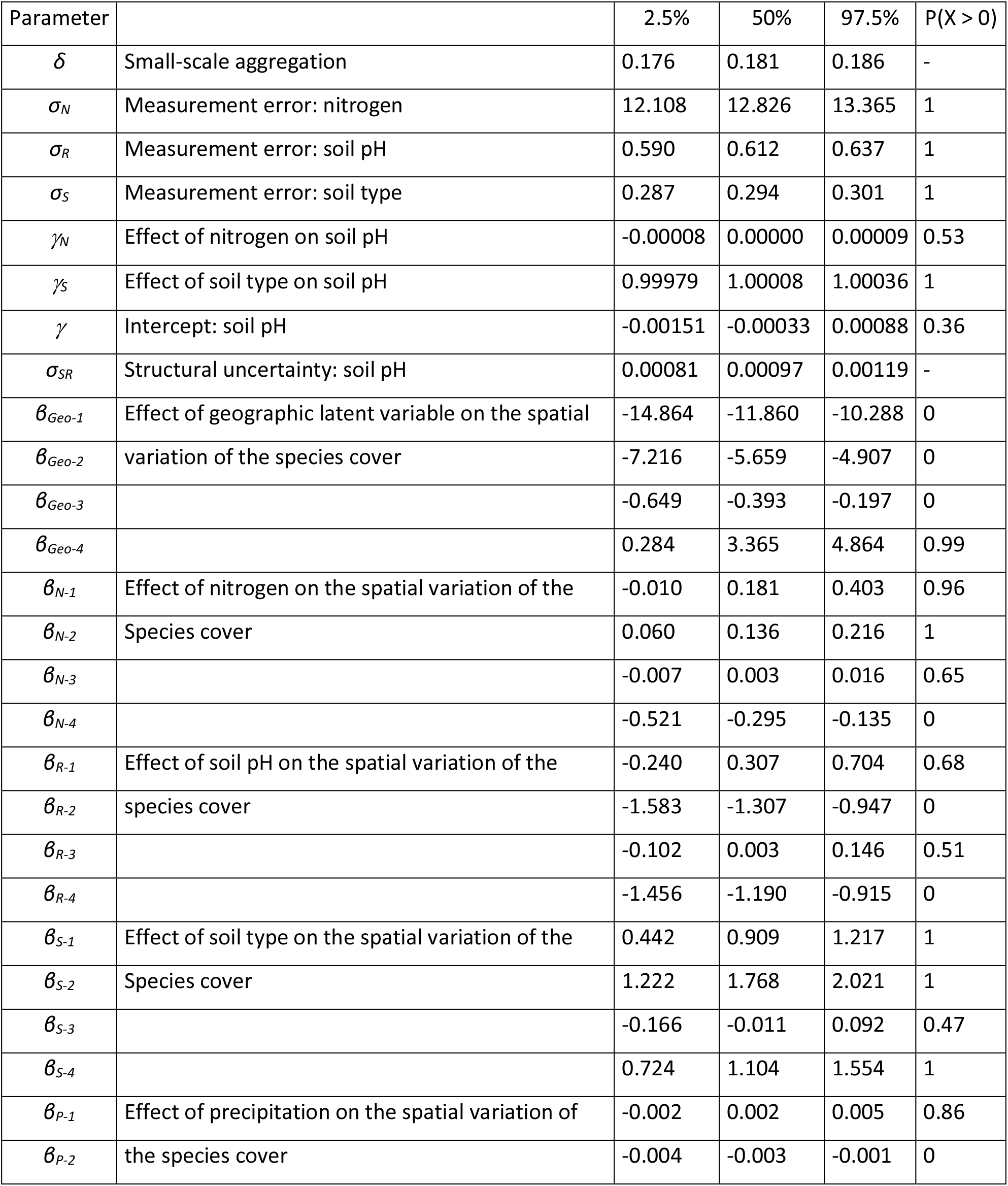
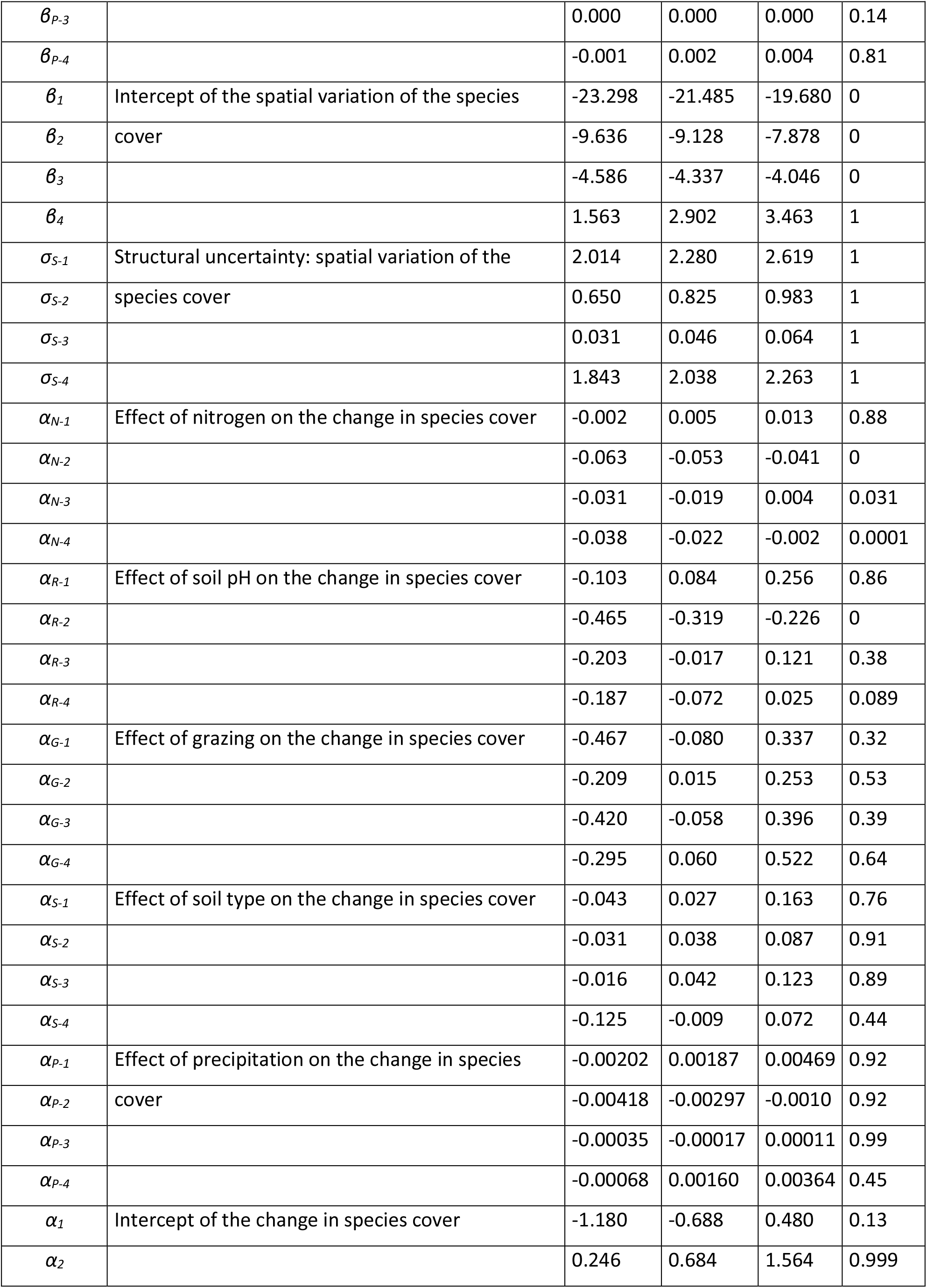
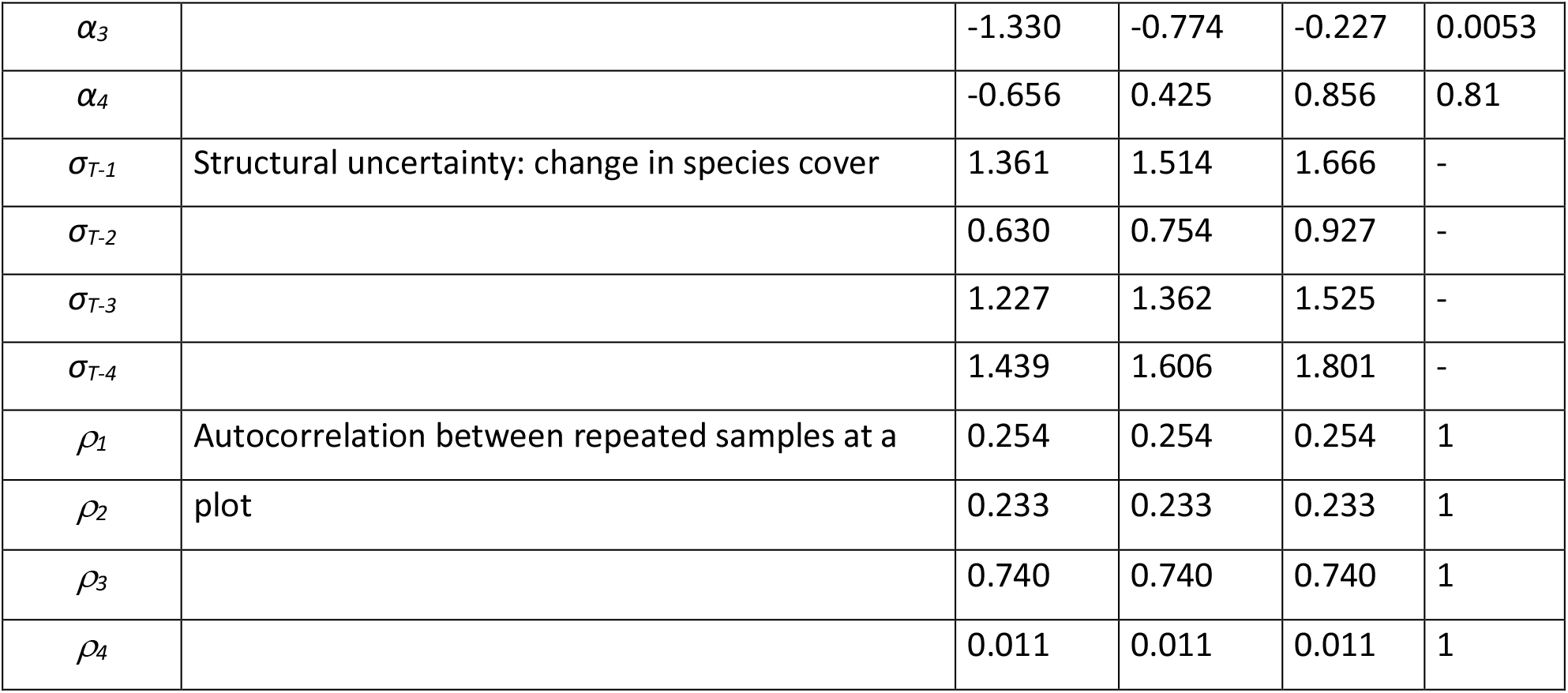
Marginal posterior distribution of SEM parameters shown by the 2.5%, 50% and 97.5% percentiles, as well as the probability that the parameter is larger than zero. The species are denoted by numbers, 1: grass, 2: other graminoids, 3: dwarf shrubs, and 4: other forbs.

Many of the regression parameters that measure the effect of the abiotic variables on the vegetation were significantly different from zero (Table 1), suggesting that the studied environmental drivers have a regulating effect on both the large-scale spatial variation in cover of the four species classes as well as local plant community dynamics in acid grasslands.

When considering the patterns of the large-scale spatial variation in cover of the four species classes, increasing nitrogen deposition had a significant positive effect on the cover of grasses and other graminoids, and a significant negative effect on the cover of other forbs. Increasing soil pH had a significant negative effect on the cover of other graminoids and other forbs. An increased clay content in the soil had a significant positive effect on the cover of grasses, other graminoids and other forbs, whereas increasing annual precipitation had a significant negative effect on the cover of other graminoids (Table 1, Fig. 6A).

**Fig. 6.**
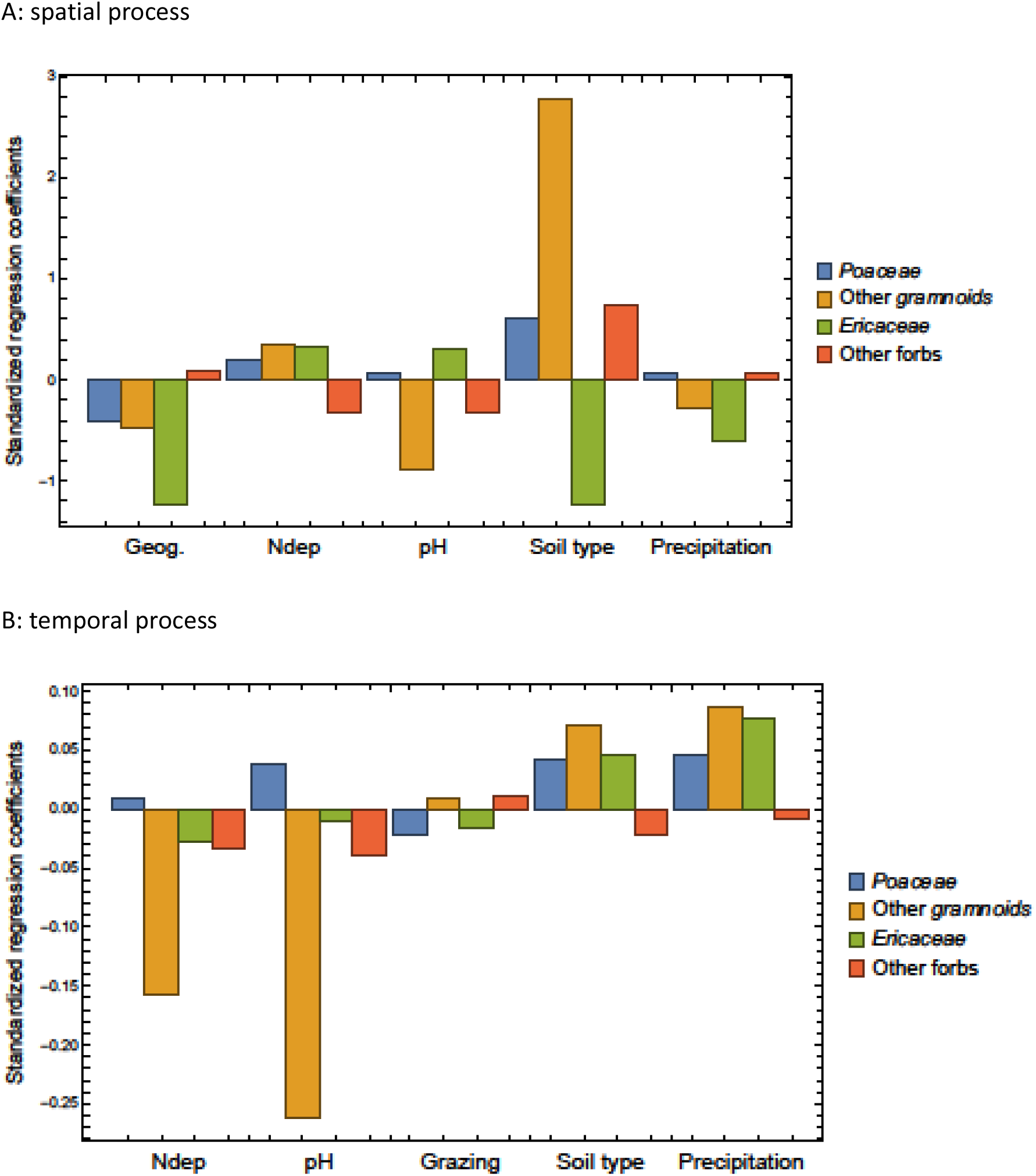
Standardized regression coefficients of the SEM for the initial spatial effects (A) and the later temporal effects (B).

There were significant regional differences in the large-scale spatial variation in cover of the four species classes that could not be explained by the studied environmental- and land-use drivers (Table 1). The regional geographic variation of the cover of grasses, other graminoids and dwarf shrubs were all positively correlated, and negatively correlated to cover of other forbs (Table 1, Fig. 6A) on the scale of approximately 50 km. (Fig 1). This finding indicates that the habitat requirements of other forbs differ from the other three species classes, and that some of the large-scale spatial variation in cover that could not be explained by the studied drivers may be explained by hitherto unexplored factors.

As mentioned above, the temporal process model explained a relatively small part of the observed variation in the change in cover during the study period. Never-the-less, increasing nitrogen deposition had a significant negative effect on the cover of other graminoids, dwarf shrubs and other forbs. Increasing soil pH had a significant negative effect on the cover of other graminoids, whereas neither soil clay content nor annual precipitation nor livestock grazing had any significant effects on the observed changes in cover (Table 1, Fig. 6B).

There were no significant effects of nitrogen deposition on soil pH (*γ_N_*, Table 1), whereas soil pH was found to be significantly higher on clayey soils compared to more sandy soils (*γ_s_*, Table 1). Based on these results, it was concluded that the effect of nitrogen deposition mainly was through direct effects on the vegetation and not by possible acidification of the soil, whereas the effect of soil type on the vegetation included both direct effects of soil type and indirect effects mediated by the effect of soil type on soil pH.

## Discussion

The study corroborates the findings of earlier studies that nitrogen deposition affects the species composition of acid grasslands (Helsen et al. 2014; Stevens et al. 2011; Stevens et al. 2006). Generally, increasing nitrogen deposition leads to more grass-dominated acid grassland habitats at the expense of the cover of forbs, both in the observed large-scale spatial variation at the onset of the study in 2007 and in the observed temporal changes in cover during the study period from 2007 to 2014. It is remarkable that relatively strong and significant effects of nitrogen deposition were found even though the range in nitrogen deposition in Denmark is relatively modest. Interestingly, nitrogen deposition was found to have a relatively strong negative effect on the changes in the cover of other graminoids and a smaller negative effect on the changes in the cover of dwarf shrubs (Fig. 6B), although there was a significant positive effect in the observed large-scale spatial cover variation for other graminoids.

Anthropogenic nitrogen deposition has reached a maximum and is currently decreasing in Denmark (Ellermann et al. 2018), but alarming changes towards more grass-dominated acid grassland habitats are still taking place, possibly as a delayed response to former nitrogen deposition levels. The estimated empirical critical load of nitrogen deposition is between 10 kg N ha^-1^ year^-1^ – 20 kg N ha^-1^ year^-1^ (Bobbink et al. 2010) compared to the mean deposition of 12.87 kg N ha^-1^ year^-1^ in Danish acid grassland sites. Consequently, to conserve biodiversity in acid grasslands, it will most likely be important to further reduce atmospheric nitrogen deposition, especially since more grass-dominated acid grassland habitats are expected to lead to cascading negative effects on pollinators and specialist insect herbivores.

It has been hypothesized that nitrogen deposition may lead to increased acidification in natural and seminatural habitats (Bobbink et al. 2010; Williams and Anderson 1999). However, in Danish acid grasslands there was no significant large-scale spatial association between nitrogen deposition and soil pH, and this observed non-significant effect of nitrogen deposition on soil pH is in concordance with earlier findings in wet heathlands (Damgaard 2019b; Damgaard et al. 2014). The potential soil acidification effects of nitrogen deposition are due either to nitrate leaching or removal of base cations from the system by nature management (Williams and Anderson 1999), and the earlier and present results may indicate that nitrate leaching or base cations removal from Danish light-open habitats are limited.

Soil type and soil acidity are important factors in determining the functional and realized niche of plant species and consequently play and important role in regulating plant community dynamics (Ellenberg 1979; Pärtel 2002). Generally, acidic soils have fewer plant species (Pärtel 2002), and a higher intra-specific aggregation of plant species has been observed in acid grasslands (Damgaard et al. 2013). Here, sandy soils were found to be relatively more acidic, and the effect of soil type on the vegetation included both direct effects of soil type and indirect effects mediated by the effect of soil type on soil pH. More specifically, increased clay content had a significant positive effect on the large-scale spatial variation in cover of grasses, other graminoids and other forbs, and increasing soil pH had a significant negative effect on the cover of other graminoids and other forbs. Furthermore, increasing soil pH had a significant negative effect on the change in cover of other graminoids.

The annual precipitation in the future climate in Denmark is predicted to increase, but with decreasing summer precipitation and longer summer drought periods (DMI 2017). Based on the present study, it is uncertain how this combination of more extreme weather will influence the future acid grassland vegetation, although we found that increasing annual precipitation had a significant negative effect on the large-scale spatial variation in cover of other graminoids. Generally, the effect of climate change in acid grasslands is still quite uncertain and must be monitored systematically in the future and with increased focus on extreme events (Galvánek and Janák 2008; Herben et al. 2003; Hopkins and Del Prado 2007).

Due in part to imperfect grazing data, there was considerable uncertainty on the estimated temporal effect of grazing on acid grasslands. As mentioned in the beginning, semi-natural acid grasslands only persist in the long run due to extensive grazing and occasional mowing (Galvánek and Janák 2008; Timmermann et al. 2015). Consequently, if better grazing data had existed, it is expected that a temporal effect of grazing would have been observed. To further investigate the effect of grazing on acid grassland vegetation, it will be necessary to conduct manipulated experiments and measure the effect of grazing on the inter-specific competitive interactions (e.g. Bullock et al. 2001)

The most important sources of measurement- and sampling uncertainties have been included in the hierarchical model structure using latent variables, which may be important in avoiding model- and prediction bias (Damgaard 2020b). Furthermore, the measurement- and sampling uncertainties are separated from the process uncertainty, which is important when using the fitted SEM for generating ecological predictions.

In the previously fitted hierarchical SEM model of wet heathland (Damgaard 2019b), the temporal model explained a higher proportion of the observed variation in the change of cover than found in this study. Wet heathlands are dominated by three species that each were modelled separately, and possibly the higher level of species aggregation, which are needed in this study of the more species-rich acid grasslands, puts an upper limit on the fitting properties of the temporal SEM. That is, when more species are aggregated into a single species class, then ecological predictions of the species class become more uncertain.

Although only a small part of the temporal variation in cover was explained by the SEM, it will still be useful to use local ecological predictions of the expected probability distribution of the different cover classes to set up local adaptive management plans (Damgaard 2020a). The chosen level of taxon generality, i.e. the aggregation of all higher species into the four species classes, may be relevant for specific applied applications. For example, when local ecological predictions are needed to develop local adaptive management plans for sites where the general conservation goal is to increase the cover of forbs and insect-pollinated plants. However, if the local nature management targets specific dominant and problematic plant species (Koerner et al. 2018; Pecháčková et al. 2010), then more detailed autecological information of the dominant plant species is more relevant.

This study is an example of the approach advocated by e.g. Ovaskainen et al. (2017), where large-scale environmental and species community data are fitted in relatively complicated statistical models in order to get more quantitative information of the ecological processes that control observed biodiversity patterns and changes. The increasing application of this approach in tight connection with more targeted studies of more specific ecological processes at local scales, e.g. inter-specific competition, will most likely advance our capacity to make credible ecological predictions in the future. Such quantitative ecological predictions are needed for making the science of ecology relevant for society in our effort to diminish the damaging effects of anthropogenic environmental changes.

## Acknowledgements

There were no conflicts of interest.

## Data availability

The used data are all in public domain, e.g. at https://naturdata.miljoeportal.dk/.

## Electronic supplements

The electronic supplements may be found at DOI 10.17605/OSF.IO/3BQ86

Table S1: Maximum likelihood estimates in the zero-inflated beta distribution of mean site cover for the 25 most common species in Danish acid grassland sites; mean site cover (μ), beta distribution shape parameter (v) and the probability that the species is absent from a site (γ). Note that because of plant overlap, total cover is larger than 1 (Ospina and Ferrari 2010).

Table S2: Correlation matrix of parameters.

Fig. S1. Dunn–Smyth residuals of the marginal observed cover data of the four species classes.

Mathematica notebook

